# Actin filament alignment causes mechanical hysteresis in cross-linked networks

**DOI:** 10.1101/2021.03.16.435707

**Authors:** Danielle R. Scheff, Steven A. Redford, Chatipat Lorpaiboon, Sayantan Majumdar, Aaron R. Dinner, Margaret L. Gardel

## Abstract

Cells dynamically control their material properties through remodeling of the actin cytoskeleton, an assembly of cross-linked networks and bundles formed from the biopolymer actin. We recently found that cross-linked networks of actin filaments reconstituted in vitro can exhibit adaptive behavior and thus serve as a model system to understand the underlying mechanisms of mechanical adaptation of the cytoskeleton. In these networks, training, in the form of applied shear stress, can induce asymmetry in the nonlinear elasticity. Here, we explore control over this mechanical hysteresis by tuning the concentration and mechanical properties of cross-linking proteins in both experimental and simulated networks. We find that this effect depends on two conditions: the initial network must exhibit nonlinear strain stiffening, and filaments in the network must be able to reorient during training. Hysteresis depends strongly and non-monotonically on cross-linker concentration, with a peak at moderate concentrations. In contrast, at low concentrations, where the network does not strain stiffen, or at high concentrations, where filaments are less able to rearrange, there is little response to training. Additionally, we investigate the effect of changing cross-linker properties and find that longer or more flexible cross-linkers enhance hysteresis. Remarkably plotting hysteresis against alignment after training yields a single curve regardless of the physical properties or concentration of the cross-linkers.

## Introduction

The mechanical properties of eukaryotic cells are, to a large degree, determined by the actin cytoskeleton [1]. Actin monomers polymerize into semi-flexible filaments with a persistence length of approximately 10 μm. The formation of space spanning networks is controlled by myriad actin binding proteins that regulate polymer assembly dynamics and cross-linking to control the local architecture of these networks [2]. To allow for cell shape change and cytoskeletal remodeling, actin networks in vivo rapidly adjust their structure and mechanics. Stress-mediated adaptation can arise from mechanotransduction pathways that dynamically regulate actin cytoskeletal composition [3]. Alternatively, the dynamic nature of cross-linking proteins can also result in adaptation in passive networks [4–6].

The structure and rheological properties of actin networks are controlled by varying the concentration and type of protein cross-linkers [7–9]. At a sufficiently high cross-link density, the network’s elastic modulus increases nonlinearly at large strains, a phenomenon known as strain stiffening. Strain stiffening is a reversible phenomenon, arising from the nonlinear increase in the spring constant of individual semi-flexible polymers as they are stretched [10].

While filament stretching dominates the mechanics of strain stiffening, filaments can also buckle under shear, creating a locally weakened region [11,12]. Changes in cross-linker concentration or filament orientation can influence the relative likelihood of shear-induced bending and stretching, impacting the mechanical response [12–15]. The physical properties of cross-linkers, such as their length and flexibility, also influences network structure and mechanics [16].

We recently showed that actin networks cross-linked with the protein filamin exhibit mechanical hysteresis [4]. In contrast to strain stiffening, mechanical hysteresis is a stress-induced and direction-dependent modification to elastic properties that is maintained long after the applied stress is removed but can then be modified by subsequent stress applications. These networks have an asymmetric response to strain, where higher stress is required to shear the network in the direction of the previously applied stress than in the reverse direction. Simulations suggest the asymmetric response results from shear induced filament alignment [4,5]. These simulations, however, did not investigate how filaments realign under stress or how the nature of the cross-linking protein might affect this reorganization. While the mechanics and affinity of the cross-linking proteins are known to play an important role in rheology, their effect on mechanical hysteresis and the adaptive properties of actin networks is unknown.

Here, we explore how cross-linker properties and concentration can be used to modify the extent of mechanical hysteresis. Furthermore, simulations allow us to directly probe the internal structure of networks and how changes correlate with hysteresis. We find that hysteresis requires networks to have a non-linear response to strain and correlates strongly with the ability of filaments to align under stress.

## Methods

### Protein Purification

Monomeric actin (G-actin) is purified from rabbit skeletal muscle acetone powder (Pel Freeze Biologicals, Product code: 41008-3) using a procedure adapted from [17]. Actinin and dictyostelium discoideum filamin (ddFLN) are each expressed in E. coli BL21-Codon Plus(DE3)-RP cells and purified using a HIS tag as described in [18]. Human filamin (FLN) is expressed in insect Sf9 cells and purified using a FLAG tag. All proteins are drop-frozen in liquid nitrogen and stored at −80 °C until use.

### Network Preparation

To prepare in vitro networks, 23.8 μM of G-actin and a varied concentration of cross-linker are added to Ca-buffer G (0.1 mM CaCl_2_, 2 mM TRIS, 0.2 mM ATP, 0.5 mM DTT, 1 mM NaN_3_, pH 8). Polymerization is initiated by adding 1/10 the final volume of 10x actin polymerization buffer (500 mM KCl, 10 mM MgCl_2_, 2 mM EGTA, 100 mM Imidazole) and mixing immediately before placement on the rheometer sample chamber. After loading the sample, the value of *G*’ and *G*” with time is measured at a frequency of 0.5 Hz and a strain of 0.05 to track network polymerization, characterized by an increase in both moduli. Each network is polymerized for 1.5 hrs, at which point both *G*’ and *G*” are constant with time. Crosslinker concentration is reported as a ratio *R_Cross–linker_* = [*cross – linker*]/[*actin*].

### Bulk Rheology

All rheological measurements are performed on an Anton-Paar MCR301 rheometer at 22° C using a 25 or 50 mm diameter plate and a 160 mm gap. A humidity chamber is used to prevent solvent evaporation. Each readout is performed 3 consecutive times. For analysis, the second time is used to avoid the impact of the initial acceleration. To find *γ_max_*, we perform the readout process to incrementally increasing *γ* on an untrained network. *γ_max_* is then the highest value of *γ* for which the network does not irreversibly weaken during repeated readout cycles. This process is repeated at different cross-linker concentrations and the lowest value is used. *γ_Max_* is measured separately for each cross-linker. During training the networks undergo plastic deformation and no longer relax back to *γ* = 0 under zero stress, but instead maintain a residual strain *γ_R_*. For subsequent readouts, we redefine zero strain such that *γ_R_* = 0.

### Simulations

Networks were simulated using the package AFINES, which has been previously described in [19] and is summarized here. AFINES takes advantage of a coarse-grained description of cytoskeletal components to efficiently simulate their networks. Specifically, actin filaments are parametrized as N+1 beads connected by N springs of length 1 μm; an additional N-1 springs are applied to the angles at the joints to limit bending and afford simulated filaments a persistence length similar to that measured experimentally. Cross-linkers are similarly modeled as springs of length *l* and stiffness *f* with beads on either end that can bind and unbind from filaments via a kinetic Monte Carlo scheme that preserves detailed balance. The motion of filaments and cross-linkers evolves according to an overdamped Langevin dynamics in two dimensions with a timestep Δ*t* of 10^-6^ s and Lees-Edwards boundary conditions [20].

To assess the rheological properties of simulated networks we perform training and readout shears analogously to the experiment. For what follows, we define *γ* = Δ*X/Y* as the unitless engineering strain, where *Y* is the box height and Δ*X* is the maximum horizontal displacement since the initiation of shear. Every △*t_s_* = 10^-5^ s, we apply a shear strain to bead *i* using

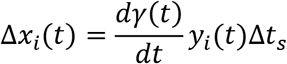

where *x_i_* and *y_i_* are the coordinates of bead *i*. We measure shear stress as (1/*A*) *dU/dγ* where *A* is the area of the simulation box and *U* is the total potential energy.

To initialize the simulations, we draw the position of the first bead of each filament or crosslinker with uniform probability within the box and introduce the remaining beads in a manner consistent with the Boltzmann distribution such as to randomly orient the crosslinker or filament. For the training, we equilibrate for 5 s and then compute *γ/dt* at every tenth step (i.e., △*t_s_* = 10^-5^ s) such as to achieve a constant shear stress *σ* using the Berendsen barostat [21]:

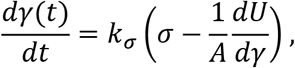

where *k_σ_* is the strength of the coupling; we substitute the resulting *γ/dt* into the equation for △*X_i_*(*t*) above, which is executed prior to the Langevin integration. As in the experiment, networks are trained to the stress that an untrained network achieves at *γ* = 0.5. Simulations are trained by applying the stress *σ* with *k_σ_* = 0.2μm/(pN s) for 10 s before allowing them to relax by approaching *σ* = 0 with *k_σ_* = 0.01 μm/(pN s) for an additional 10 s before readout is performed.

For readout simulations, we equilibrate for 5 s before applying a triangle wave of the form

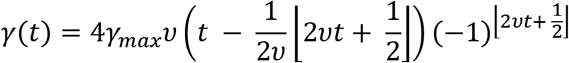

where *γ_max_* is the amplitude of the wave and *ν* is the frequency of the oscillation. For all of the simulations considered herein *γ_max_* = 1 and *ν* = 0.5 s^-1^. We measure a number of shear cycles to ensure that the simulation is stable over time. Specifically, we apply readout shear for 11 seconds such that untrained simulations have a final time, *t_final_*, of 16 s.

## Results and Discussion

### Networks’ Response to Training Stress

To investigate its origin and controlling factors, we probe mechanical hysteresis in actin networks formed with cross-linkers with various physical properties and concentrations. As such, we require a standardized method of imparting and measuring hysteresis that can be applied uniformly to all cross-linker conditions. We adapt the process developed by Majumdar et al., in which the network’s directional response to shear is measured before and after a training stress [4]. Specifically, we train the networks by subjecting them to a constant stress, *σ*, for 300 s, before allowing them to relax at *σ* = 0 for an additional 100 s (Fig. 1a). Here, training always occurs in the same direction, corresponding to clockwise rotational shear and defined as positive strain, whereas counterclockwise rotation is defined as negative strain. We measure the effect of training using a readout process, in which the network is cyclically sheared with an amplitude of *γ_max_* (Fig. 1b). Comparing readout curves taken before and after training allows us to ascertain the effect that training has on the rheological properties of cross-linked networks and reveals any asymmetric responses arising from the direction of the training stress. Since networks irreversibly break at sufficiently large strains, we set *γ_max_* to be the maximum strain at which the network can be repeatably sheared with no observed changes in its stress response, as described in methods. *γ_max_* was determined separately for each cross-linker used. The training stress for each network is individually determined as the stress required to shear the untrained network to *γ* = *γ_max_/2*.

**Figure 1:**
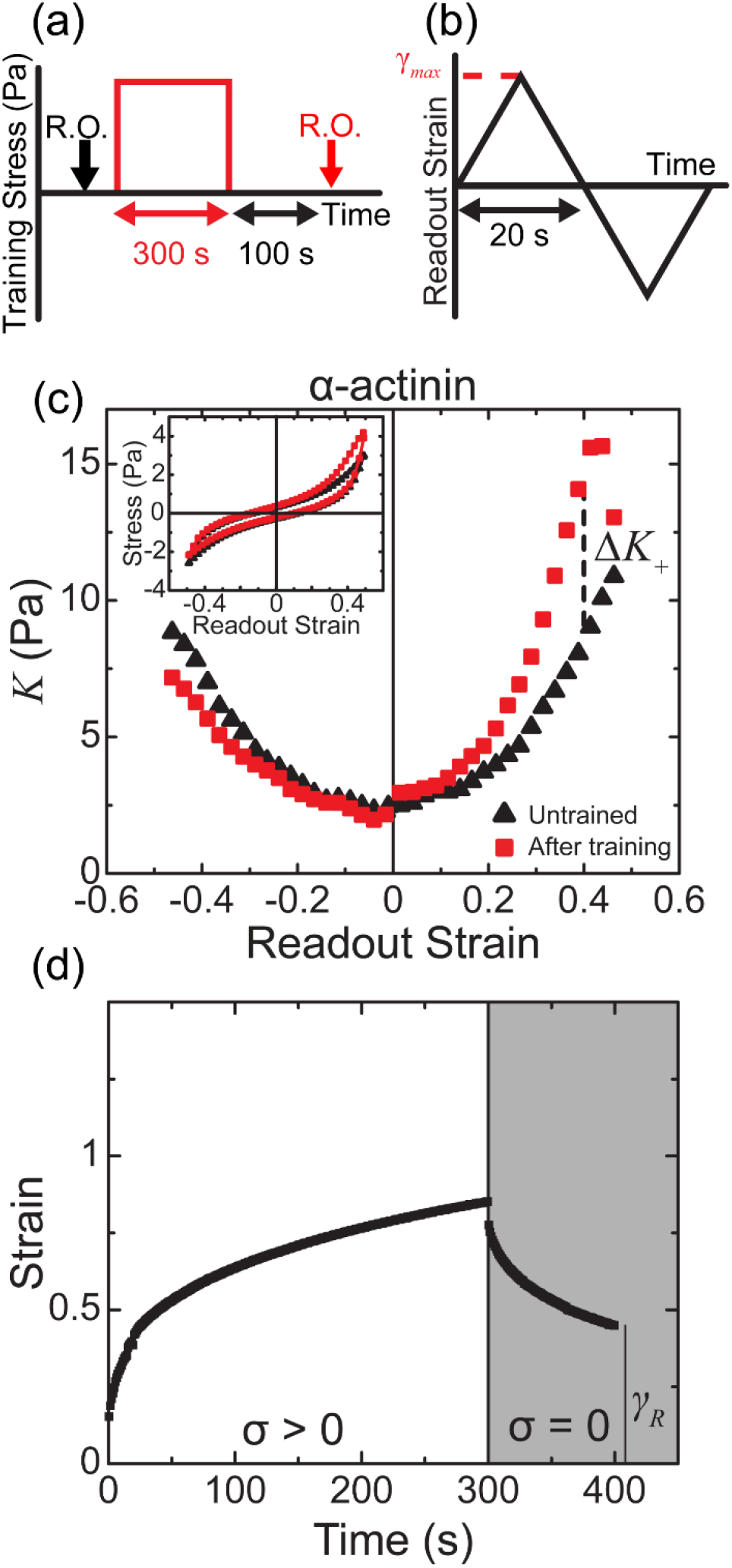
Cross-linked actin networks have asymmetric response to strain after a training stress. (a) Cross-linked actin networks are trained by applying a constant stress for 300 s before being allowed to relax at zero stress for an additional 100 s. Before and after training, the non-linear response to strain is measured using a readout process (b), in which the network is sheared in both the direction of training and the opposite direction. (c) The differential modulus *K*, measured at different strains during readout, before (black triangles) and after (red squares) training. △*K*_+_ measures the rescaled difference in the trained versus untrained value of *K. Inset*: The corresponding stress from which *K* was measured. Data shown for a network with *R_α_* = 1%. (d) Example creep response of the strain to a training stress (white region). After training (gray region), the network does not relax to its original position, leaving a residual strain *γ_R_*.

First, we measure the effect of training on networks cross-linked with α-actinin, a rigid protein able to form transient bonds between filaments, and whose rheological properties are well studied [22]. We characterize the readout response through both the stress and the differential modulus *K = dσ/dγ* as a function of the strain, *γ*. *K* can be understood as a strain-dependent elastic modulus, and any increase reflects a nonlinear, strain stiffening response. Initially, both *σ* and *K* are symmetric across *γ* = 0, as expected for an isotropic material (Fig. 1c, black triangles) [4]. We call this initial case the “untrained” network. In contrast, the networks develop an asymmetric response to shear direction after training. After training, *K* increases for *γ* > 0 and decreases for *γ* < 0 relative to the untrained case (Fig. 1c, red squares). Training thus induces a direction dependent mechanical response. We refer to this phenomenon as mechanical hysteresis and quantify it by the parameter

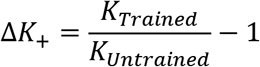

where *K_Untrained_* and *K_Trained_* are the differential moduli measured at *γ* = 0.8 * *γ_max_* before and after training, respectively. △*K*_+_ corresponds to the fractional increase in differential modulus for *γ* > 0. Furthermore, the network does not relax back to its original state after training. Instead, it undergoes a plastic deformation and maintains a residual strain, *γ_R_* (Fig. 1d). In previous work, we found that increased *γ_R_* corresponds to larger amounts of hysteresis [4]. This correlation hints at underlying changes in the network during training that could explain the observed phenomenon. However, while this observation describes the bulk phenomena of mechanical hysteresis, it does not provide any microscopic origin.

To grasp the microscopic effect that training has on cross-linked networks we turn to coarse-grained simulations using the simulation package AFINES [19]. Past modeling efforts initialized simulations in putative post-training geometries to evaluate plausible mechanisms of mechanical hysteresis, namely that the effect can be explained by post-training nematic alignment of filaments [4]. However, here we endeavor to explicitly simulate the training process to gain a more complete understanding of shear-induced rearrangements. The model and simulations are described in Methods. In brief, we represent networks in two dimensions by bead-spring filaments that are connected by cross-linkers with length *l* and stiffness *f* that can bind and unbind filaments with rates *k_on_* and *k_off_* (Fig. 2a). Our simulations specifically contain 500 7-μm filaments initialized randomly in a 20 x 20 μm box with periodic boundary conditions. Training and readout follow protocols analogous to the experiments as described in Methods.

**Figure 2:**
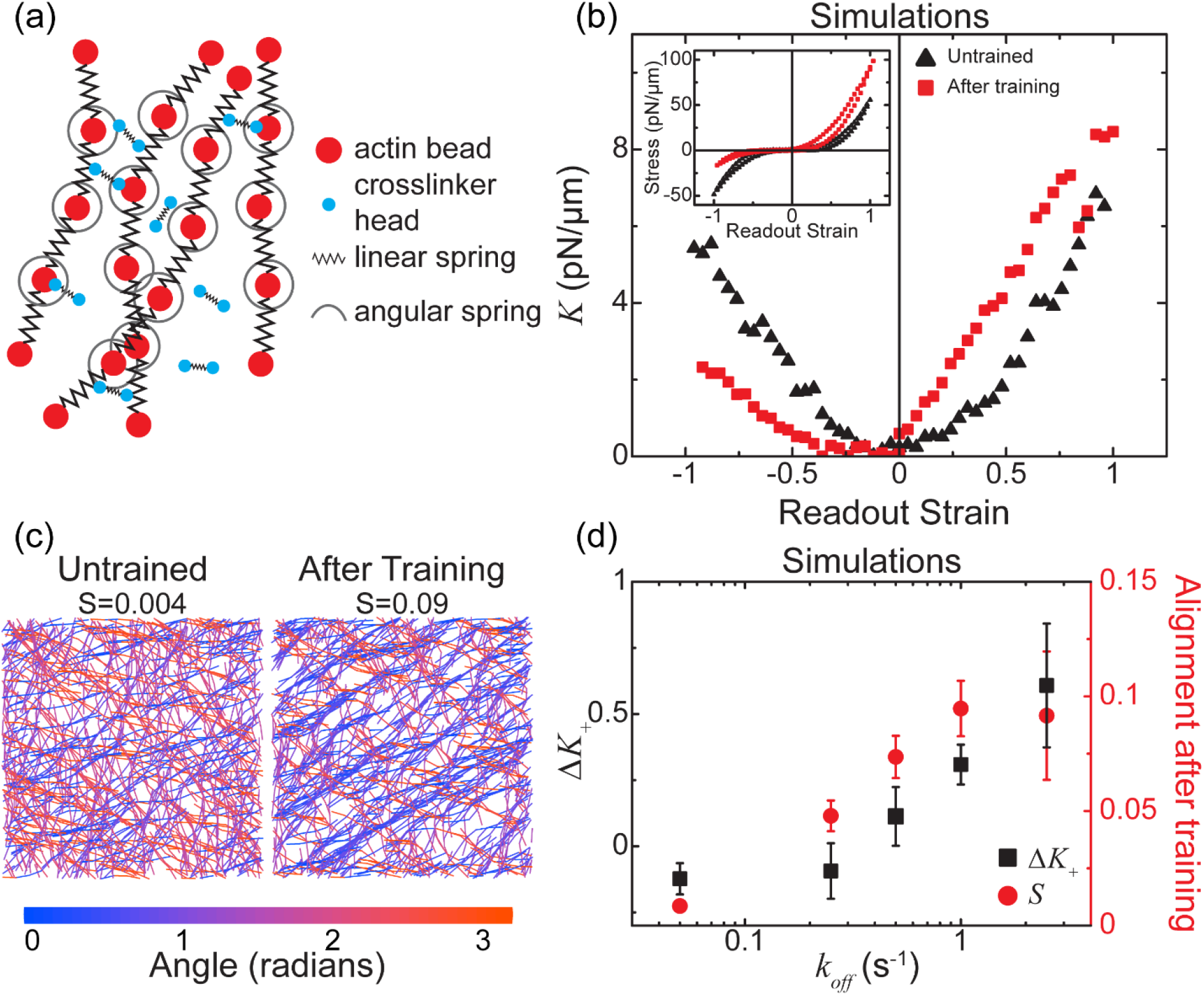
Simulated networks develop mechanical hysteresis due to alignment of filaments during training. (a) Schematic of the AFINES model. Filaments and cross-linkers are parametrized as beads connected by springs with cross-linkers able to bind and unbind according to a kinetic Monte Carlo scheme. (b) The differential modulus *K*, as measured in simulations before (black triangles) and after (red squares) training. *Inset*: The corresponding stress during readout. Data from simulations with *k_off_* = 0.5 s^-1^. (c) Images of simulated network after equilibration but before the onset of training (left) and after training immediately before readout (right). Color corresponds to filament orientation. (d) △*K*_+_ (black squares) and filament alignment *S* (red circles) in simulated networks with varying cross-linker off rate. All simulations shown have *R_Sim_* = 9 μm^-2^, *l* = 0.15 μm, and *f* = 100 pN/μm. Error bars are standard deviation of 5 independent simulations.

We observe a mechanical hysteresis response similar to that seen in experiment with a cross-linker density of 9 μm^-2^, *l* = 0.15 μm, *f* = 100 pN/μm, *k_off_* = 0.5 s^-1^, and a *k_on_ /k_off_* = 10 (Fig. 2b). Specifically, the trained networks in simulation stiffen in the direction of training and soften in the opposite direction (Fig. 1f). Having successfully recapitulated mechanical hysteresis with a heretofore uninvestigated cross-linker and a simulation that explicitly models training we now turn to investigate the origins of the phenomenon.

### Measuring Filament Alignment

Majumdar and colleagues’ past investigation of mechanical hysteresis in similar networks suggested that the ability of the network to rearrange in response to training is the origin of the mechanical hysteresis response. To investigate whether such rearrangements occur in our simulations, we measure the nematic order parameter

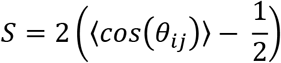

where *θ_ij_* is the angle between two filament sections. For the simulated network in Fig. 2b, after equilibration but before the onset of training, the network is largely isotropic with *S* =0.004 (Fig. 2c). In contrast, the filaments in the trained network before readout are aligned with *S* =0.09, over an order of magnitude increase. These results indicate that networks do indeed rearrange in response to the training stress.

We test the importance of these network structural changes in developing hysteresis by altering the unbinding rate of cross-linkers in simulations. If alignment of filaments is indeed how mechanical hysteresis develops, then cross-linkers with lower unbinding rates will lead to less structural rearrangement and lower values of △*K*_+_. To test this hypothesis, we first simulate a network with a crosslinker density, *R_Sim_*, of 9 μm^-2^ and an extremely slow off rate of *k_off_* = 0.05 s^-1^ and *k_on_ /k_off_* = 100 (Fig. 2d). In this low rearrangement regime, we find that the network does not exhibit the mechanical hysteresis response following training. Furthermore, when we allow *k_off_* to vary from 0.05 s^-1^ to 3 s^-1^ while keeping the ratio of *k_on_ /k_off_* = 10, we find that decreasing off rate leads to reduced △*K*_+_ (Fig. 2d). As *k_off_* drops below 0.25 s^-1^ the network does not respond to training at all. Furthermore, these changes in △*K*_+_ correspond to changes in alignment, which similarly increases with *k_off_*, confirming that the transient nature of cross-linkers is essential for filament rearrangements. Changes in network structure during training are thus vital for mechanical hysteresis, and sufficiently reducing these rearrangements eliminates hysteresis.

### Dependence on Cross-linker Concentration

We next ask how altering the architecture of the network influences hysteresis by varying the cross-linker concentration, which impacts both network structure and rheology [8,15]. We begin by training experimental networks cross-linked with various concentrations of α-actinin, *R_α_*, and observe a drastic change in the hysteresis. For example, Fig. 3a shows a typical readout for a network with *R_α_* = 5%, which responds differently to training than the previously shown network with lower *R_α_*. Here, training has a minimal effect on the network for strains *γ* > 0, where the readout values of both stress and *K* do not change with training. In contrast, the differential modulus decreases drastically after training for *γ* < 0. Training can thus have a large effect in one direction, here *γ* < 0, while having almost no impact in the opposite one. These changes contrast with the previously shown network with *R_α_* = 1%, where training affects *K* across all strains (Fig 1c). In this case, the effect of training is about equal in both directions. We thus observe asymmetry stemming from two independent responses. Training can make the network stiffer and increase *K* when *γ* > 0, and it can soften the network and decrease *K* when *γ* <0. While the latter is interesting and is discussed further in the conclusions, here we focus on *γ* > 0, where *K* increases after training.

We further measure changes in *K* over additional cross-linker concentrations, specifically between *R_α_* = 0.05 and 10 molar percent relative to actin. Remarkably, △*K*_+_ has a non-monotonic dependence on cross-linker concentration. While increasing cross-linker concentration initially corresponds to an increase in △*K*_+_, for *R_α_* > 1% the degree of mechanical hysteresis actually decreases (Fig. 3b). Maximum hysteresis is thus achieved at *R_α_* ≈ 1%. We can understand the initial increase in △*K*_+_ with concentration by comparing it to the onset of strain stiffening. At low concentration of *R_α_* ≤ 0.2% where △*K*_+_ is close to zero (Fig. 3b, gray region), networks have a linear response to shear. Fig. 3c shows a typical example at *R_α_* = 0.2%, where *K* is relatively constant regardless of strain. △*K*_+_ begins to increase at *R_α_* > 0.2%, which is importantly also the onset of a non-linear shear response. At these concentrations *K* increases with the magnitude of strain, as shown in Fig. 3d which depicts a typical example at *R_α_* = 0.5%. The correspondence between strain stiffening and the increase in △*K*_+_ is consistent with previous work, which found that similar non-linearity was required for asymmetric response to strain in simulated actin networks [12]. It therefore makes sense that the onset of strain stiffening and hysteresis occur at the same concentration. While nonlinear shear response explains the increase in hysteresis at small cross-linker concentrations, to understand why it decreases at higher concentrations we turn to simulations.

**Figure 3:**
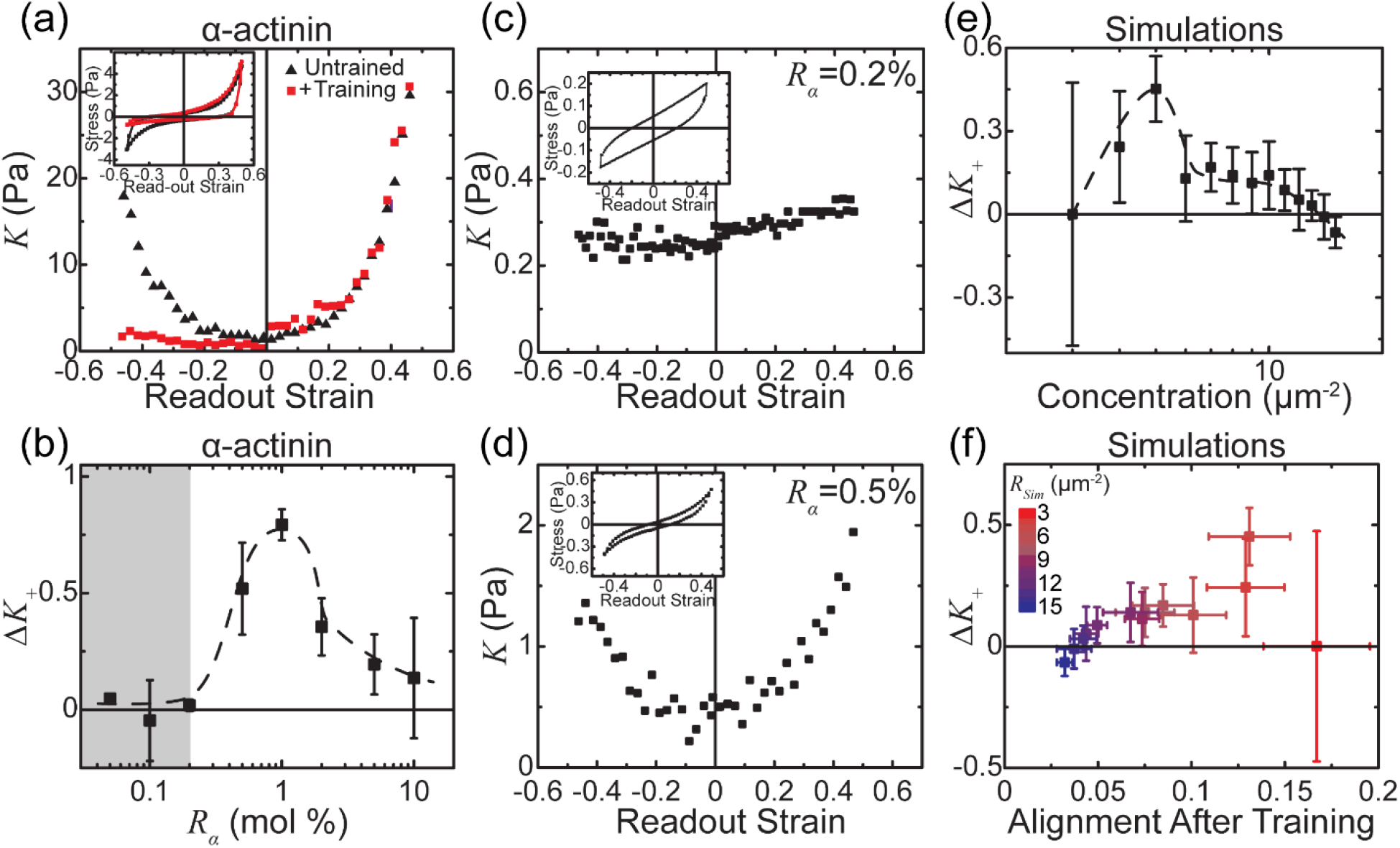
Hysteresis varies with cross-linker concentration. (a) *K* as a function of strain for a network with *R_α_* = 0.05% before (black triangles) and after (red squares) training. *Inset*: The corresponding stress vs. strain. (b) Hysteresis, characterized by △*K*_+_, plotted as a function of α-actinin concentration. △*K*_+_ is small at low and high concentrations with a peak around *R_α_* = 1%. Gray region represents concentrations networks that do not strain stiffen. (c,d) The differential modulus *K* during readout of typical untrained networks with (c) *R_α_* = 0.2%, where the network has a linear response to strain, or (d) *R_α_* = 0.5%, where the network strain stiffens. *Inset*: the corresponding stress measured during readout. (e) △*K*_+_ for different cross-linker concentrations in simulated networks. (f) More alignment of filaments after training correlates with increased △*K*_+_ in simulations across all cross-linker concentrations that permit strain stiffening. Data is for *l* = 0.15 μm, *f* = 100 pN/μm, and *k_off_* = 0.5 s^-1^. Error bars in (b), (e), and (f) are standard deviation of at least two independent experimental samples or five independent simulations. Dotted lines in (b,e) are to guide the eye.

In simulations, we can directly probe the microscopic structure of networks to investigate why mechanical hysteresis decreases at high cross-linker density. First, we perform simulations from cross-linker concentrations of 3 μm^-2^ to 15 μm^-2^ to ensure the results are consistent with experiments. Indeed, in these simulations △*K*_+_ exhibits a non-monotonic dependence on cross-linker concentration (Fig. 3e). As shown in Fig. 2c, in simulation, training is related to an increase in network alignment, so we ask whether such an effect could explain the non-monotonicity. As before, we interrogate the alignment of the network after training but before readout. We find that when we plot alignment after training against △*K*_+_ for each trial, a strong correlation emerges (Fig. 3f). This plot indicates that while at low cross-linker concentrations the network is able to rearrange in response to training, at higher concentrations, the greater number of cross-linkers actually prevents such rearrangements and thus precludes the trained network from exhibiting a strong increase in *K* relative to untrained networks. The sole outlier occurs at the lowest cross-linker concentration, where the density of cross-linkers is small enough to allow rearrangement without a corresponding increase in hysteresis. No matter how much they align under training stress, strain weakening networks do not exhibit mechanical hysteresis. Adjusting cross-linker density is one way to tune the mechanical hysteresis of a network by controlling both shear stiffening and filaments’ ability to rearrange under stress. We now ask whether there are concentration independent mechanisms of tuning the response.

### Impact of Cross-linker Properties

Thus far we have considered only α-actinin as our experimental cross-linker, but we know that networks connected by various cross-linkers can exhibit unique mechanical properties. As such, we now investigate how the physical properties of the cross-linkers in these networks affects mechanical hysteresis. Specifically, we compare networks cross-linked by α-actinin to ones cross-linked with two variants of filamin: from Dictyostelium discoideum (ddFLN) and from humans (FLN) (Fig. 4a). These proteins allow us to study the importance of cross-linker length and flexibility on mechanical hysteresis. While, at 40 nm, ddFLN is about the same length as α-actinin, it is much more flexible [23,24] and therefore allows us to alter flexibility without changing length. FLN is similarly flexible, but with a contour length of approximately 160 nm [25]. Because it is much longer than the other two cross-linkers, it can be used to measure the impact of cross-linker length. These physical differences greatly impact the rheological properties of untrained networks, leading us to investigate their effect on mechanical hysteresis.

**Figure 4:**
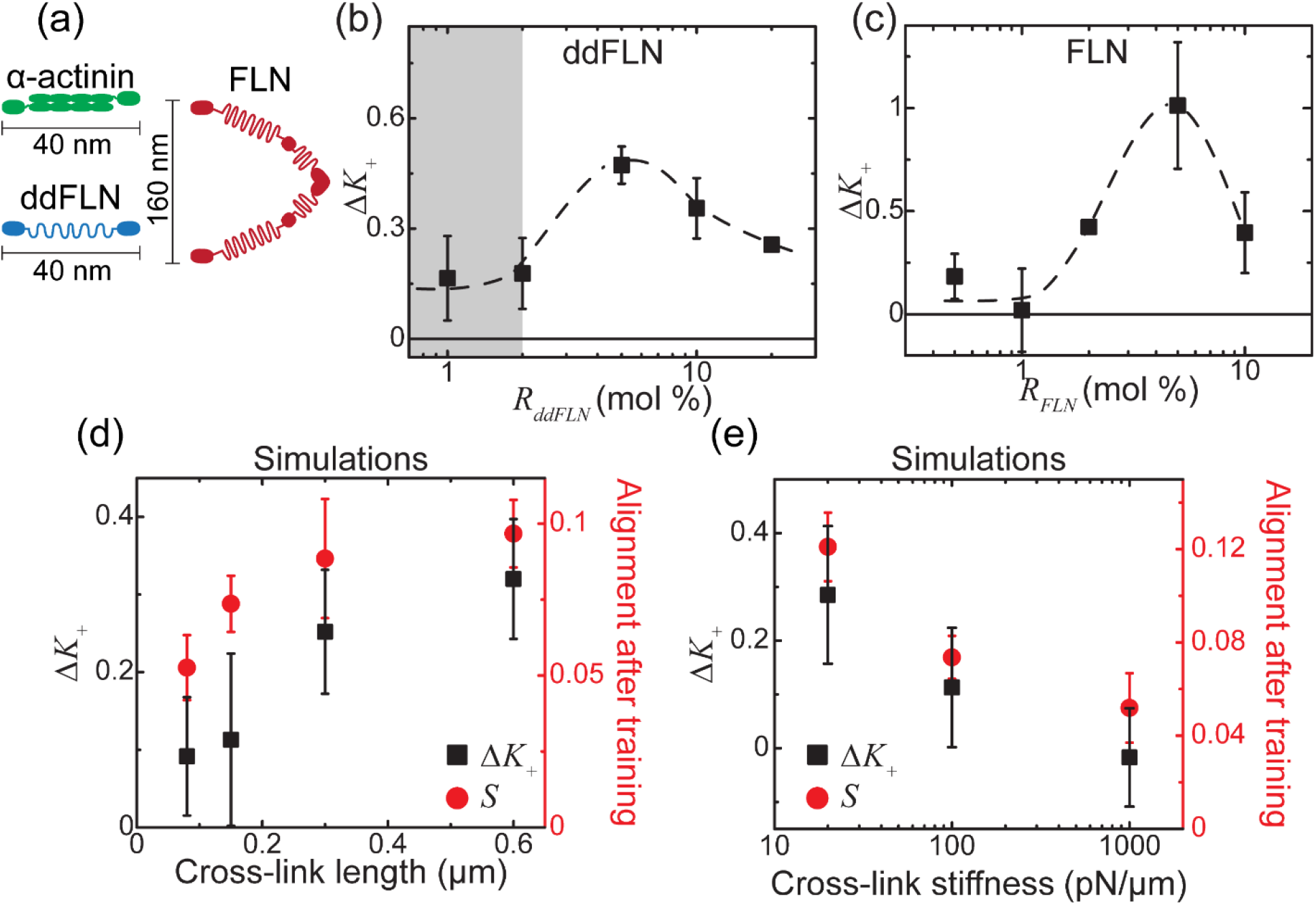
Hysteresis persists across a range of cross-linker properties. (a) Cartoon depiction of the different cross-linkers used. (b,c) △*K*_+_ at varying concentrations of (b) ddFLN or (c) FLN. Dotted lines are to guide the eye. (d,e) △*K*_+_ (black squares) and nematic order parameter *S* (red circles) in simulated networks with varying cross-linker (d) length, (e) stiffness. Simulations are with *R_Sim_ = 9* μm^-2^, *l* = 0.15 μm, *f* = 100 pN/um, and *k_off_* = 0.5 s^-1^, unless otherwise noted. Error bars in each panel are standard deviation of at least two independent experimental samples or five independent simulations.

We find that, while △*K*_+_ has different values and peaks at various concentrations depending on cross-linker, the non-monotonic dependence of hysteresis on cross-linker concentration is robust across all cross-linkers used. First, changing cross-linkers does not eliminate hysteresis, and networks cross-linked with ddFLN also develop hysteresis in response to training. Furthermore, △*K*_+_, which increases at low cross-linker concentrations before peaking and subsequently decreasing at higher concentrations, has similar dependence on concentration as for networks cross-linked with α-actinin (Fig 4b). Despite the qualitative similarities, for networks cross-linked with ddFLN, the maximum hysteresis achieved, △*K*_+_ = 0.47 ± 0.05, is smaller than that of α-actinin networks, which have a maximum △*K*_+_ = 0.79 ± 0.05. Additionally, these maxima occur at different concentrations. Whereas in α-actinin networks △*K*_+_ peaks at *R_α_* ≈ 1%, ddFLN networks show a peak △*K*_+_ at *R_ddFLN_* ≈ 5%. Notably, though it occurs at a different concentration, the initial increase in △*K_+_* in ddFLN networks still corresponds to the onset of strain stiffening (Fig. S1, ESI). Similarly, hysteresis in human FLN networks has a non-monotonic concentration dependence, but with a higher peak value (Fig. 4c). As with ddFLN, the concentration dependence of △*K*_+_ in FLN networks peaks at *R_FLN_* ≈ 5%. However, with △*K*_+_ = 1.01 ± 0.31, the maximum response of FLN networks is about twice as high as of those cross-linked with ddFLN. These cross-linkers follow the same trend observed in α-actinin networks, where hysteresis depends non-monotonically on cross-linker concentration. On the other hand, cross-linker properties do affect the magnitude of hysteresis. Because natural cross-linkers vary along a number of axes simultaneously, it is difficult to untangle how each physical difference contributes to the change in response. As such we turn again to simulation, where we are able to alter cross-linker properties independently.

Inspired by the difference between FLN and the other two cross-linkers, we examine how cross-linker length and flexibility impact mechanical hysteresis. In these simulations we fix *k_Off_* = 0.5 s^-1^ and *R_Sim_* = 9 μm^-2^, a concentration at which there is still some hysteresis but where the high concentration precludes most of the training induced alignment seen at the peak. As we vary cross-linker length from *l* = 0.08 μm to *l* = 0.6 μm, we find a corresponding increase in the magnitude of △*K*_+_ from △*K*_+_ = 0.09 ± 0.08 to △*K*_+_ = 0.32 ± 0.08 (Fig. 4d). This increase is accompanied by an increase in filament alignment. Another potentially important difference between experimental cross-linkers is their stiffness. We vary stiffness of *l* = 0.15 μm cross-linkers from *f* = 20 pN/μm to *f* = 1000 pN/μm and find that both △*K*_+_ and *S* decrease with increasing stiffness (Fig. 4e). Networks with longer or more flexible cross-linkers are thus more able to rearrange during training, leading to increased hysteresis, similar to the effect seen at high *k_off_* or cross-linker concentrations. It makes sense that short or rigid cross-linkers constrain the rearrangement of actin filaments more than long or flexible cross-linkers. We thus look at how well alignment predicts hysteresis.

When we varied cross-linker concentration and properties we found that the changes in hysteresis correlated with alterations in network alignment after training, leading us to ask how well alignment explains hysteresis across all conditions. Remarkably, when we plot alignment against △*K*_+_, changes due to altering different cross-linker physical properties or concentration collapse to a single line (Fig. 5). This trend suggests that the ability of the constituent filaments to rearrange under training stress is the main determinant of △*K*_+_ in these networks. We thus find two conditions necessary for mechanical hysteresis: the network needs to strain stiffen, and filaments must be able to rearrange under stress.

**Figure 5:**
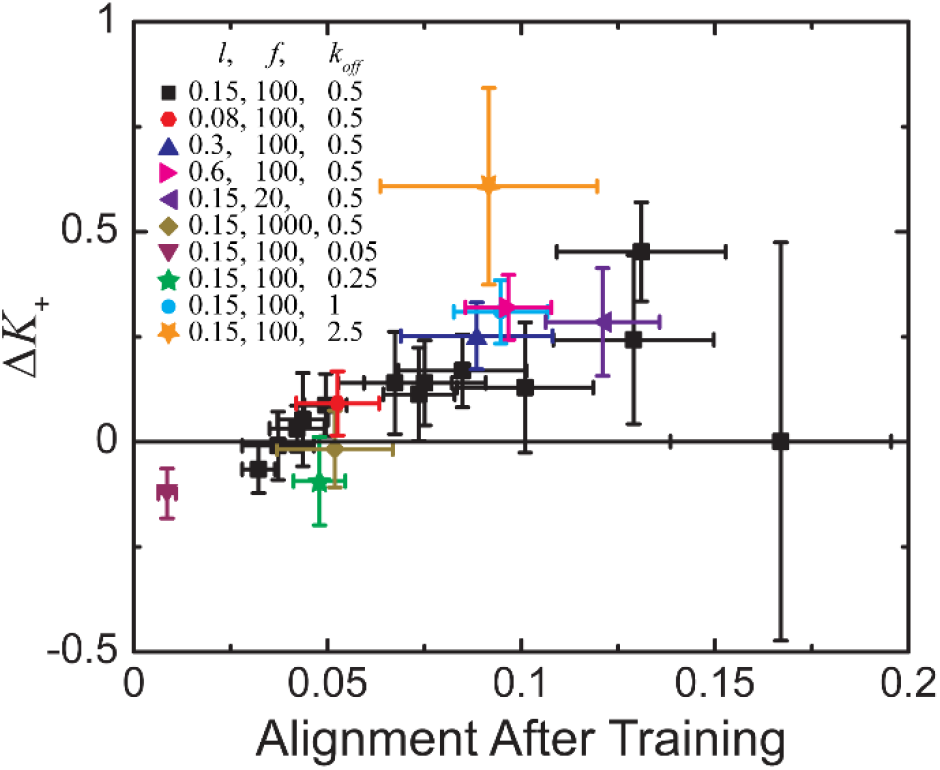
△*K*_+_ increases with filament alignment across all cross-linker parameters in simulations. All data with *R_Sim_* = 9 μm^-2^, except black squares which represent a variety of concentrations as shown in Fig 3f. Error bars are standard deviation of five independent simulations.

## Conclusion

We have shown that cross-linked actin networks display mechanical hysteresis in the form of a direction-dependent response to shear after the application of a training stress. This asymmetry results from an increase in *K* compared to the untrained network for *γ* > 0, and a decrease in *K* for *γ* < 0. We find that this increase, characterized by △*K*_+_, changes non-monotonically with cross-linker concentration. Importantly, △*K*_+_ begins to increase at the same concentration as the onset of strain stiffening. Using simulations where we can directly observe internal structural changes, we find that the decrease in △*K*_+_ at high concentrations corresponds to a drop in filament alignment after training. Furthermore, increasing the unbinding rate, flexibility, or length of cross-linkers also increases alignment, leading to higher values of △*K*_+_. This form of mechanical hysteresis thus depends on two conditions: the network having a nonlinear response to strain and the ability of the constituent filaments to rearrange under stress.

While alignment describes the increase in *K* for *γ* > 0, we also observe a second manifestation of mechanical hysteresis that is not so readily explained. In addition to its increase at positive strains, we observe that training leads to a decrease in *K* for *γ* < 0. This effect increases with cross-linker density and becomes especially pronounced at high concentrations even as filament alignment and △*K*_+_ decrease. It is thus possible to tune not only the degree of hysteresis observed but also its structure. Our simulations, however, are unable to replicate the effect of cross-linker concentration on *K* for *γ* <0. Instead, in simulations the decrease in *K* at these strains is remarkably consistent regardless of the concentration or physical properties of the cross-linkers. This discrepancy could be due to the lack of explicit hydrodynamics or the lack of filament entanglement, but further research will be necessary to determine how these might affect *K* for *γ* < 0.

This study demonstrates a hysteresis response in cross-linked polymers that can be tuned by adjusting cross-linker properties and concentration. It can therefore inform the creation of materials that passively adapt to stress. It additionally suggests ways that the degree of mechanical hysteresis can be tuned through changes in the concentration and physical properties of the cross-linkers. While we have demonstrated several knobs that can be tuned to change this response, these results suggest that any other method of altering the amount of achievable alignment should also allow for the tuning of mechanical hysteresis. Future studies could look at additional ways to tune these responses. For example, it is possible that other factors such as actin turnover rate or length could affect the ability of filaments to both align under stress and maintain this alignment over time, allowing the creation of materials that adapt more quickly to stress. Additionally, these factors could tune how quickly networks relax back to their untrained state, leading to mechanical hysteresis with shorter or longer lifetimes. It might also be possible to encode multiple hysteresis responses by training the network in an orthogonal direction. These studies would provide a greater understanding of how hysteresis responses can arise in both biological and artificial cross-linked networks.

