## Supplemental Figure 1 for "Actin filament alignment causes mechanical hysteresis in cross-linked networks"

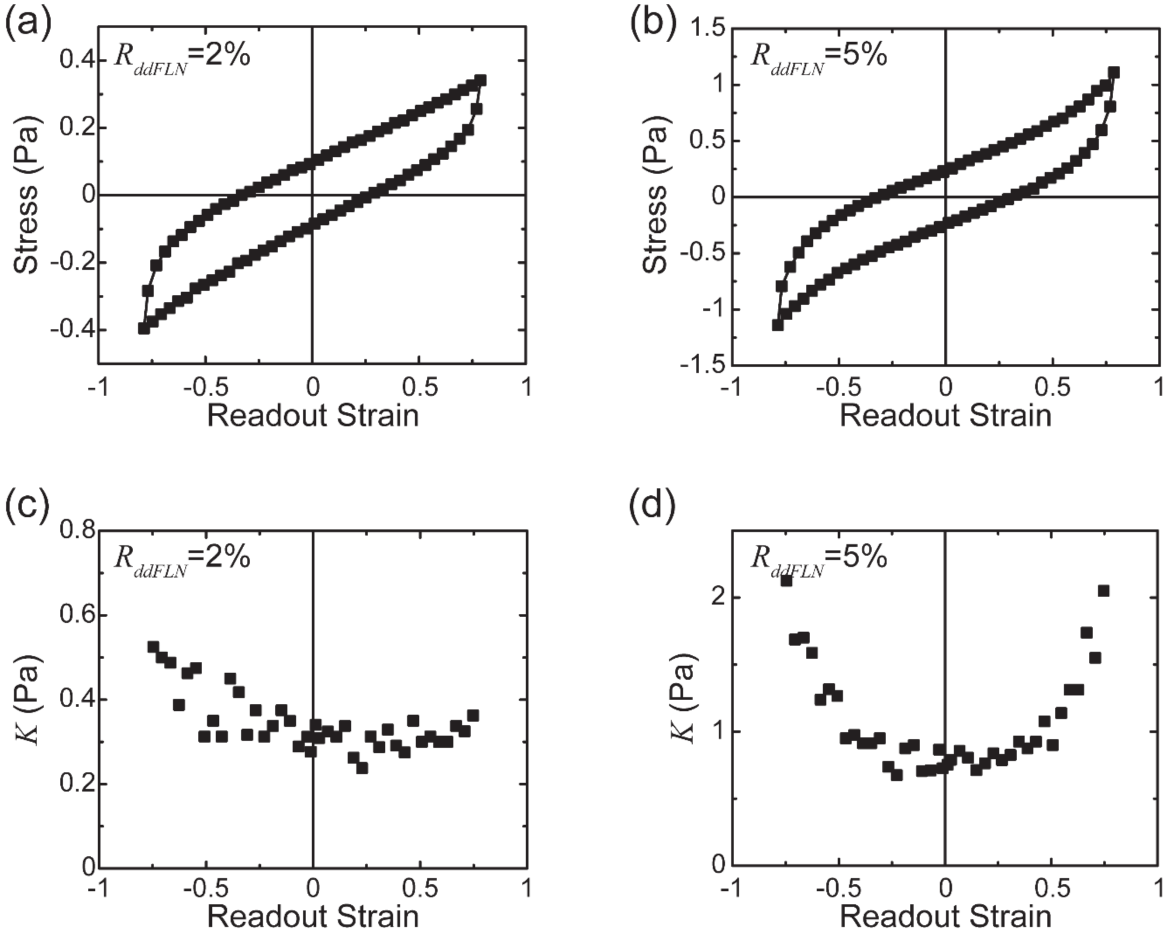


Figure S1: Strain stiffening in ddFLN networks. (a,b) Stress during readout for untrained ddFLN networks with (a) $R_{ddFLN} = 2$ or (b) $R_{ddFLN} = 5\%$, concentrations at which the network has a linear or nonlinear response to strain, respectively. (c,d) The corresponding differential modulus $K$ for networks with (c) $R_{ddFLN} = 2\%$ or (d) $R_{ddFLN} = 5\%$.
